# The cervical microbiota of Hispanics living in Puerto Rico is highly volatile and dominated by *Lactobacillus iners* regardless of HPV status

**DOI:** 10.1101/2023.05.05.539619

**Authors:** Daniela Vargas-Robles, Josefina Romaguera, Ian Alvarado-Velez, Eduardo Tosado-Rodríguez, Anelisse Dominicci-Maura, Maria Sanchez, Kara J. Wiggin, Jack A. Gilbert, Magaly Martinez-Ferrer, Larry J. Forney, Filipa Godoy-Vitorino

## Abstract

**Introduction:** The cervicovaginal microbiota is influenced by host physiology, immunology, lifestyle, and ethnicity. We hypothesized that there would be differences in the cervicovaginal microbiota among pregnant, non-pregnant and menopausal women living in Puerto Rico with and without Human Papillomavirus (HPV) infection and cervical cancer. We specifically wanted to determine if the microbiota associated with variation in cervical cytology. A total of 294 women comprised of reproductive-age non-pregnant (N=196), pregnant (N=37), and menopausal (N=61) women were enrolled. The cervicovaginal bacteria was characterized by 16S rRNA amplicon sequencing, the HPV were genotyped with SPF10-LiPA, and cervical cytology was quantified. High-risk HPV (HR-HPV, 67.3%) was prevalent, including genotypes not covered by the 9vt HPV vaccine. Cervical lesions (34%) were also common. The cervical microbiota was dominated by *Lactobacillus iners*. Pregnant women in the 2^nd^ and 3^rd^ trimesters had decreased diversity and a decreased abundance of microbes associated with bacterial vaginosis. Women in menopause had greater alpha diversity, a greater proportion of facultative and strictly anaerobic bacteria, and higher cervicovaginal pH than pre-menopausal women. Cervical lesions were associated with greater alpha diversity. However, no significant associations between the microbiota and HPV infection (HR or LR-HPV types) were found. The cervicovaginal microbiota women living in Puerto Rican were either dominated by *L. iners* or diverse microbial communities regardless of a woman’s physiological stage. We postulate that the microbiota and the high prevalence of HR-HPV, increase the risk of cervical lesions of women living in Puerto Rico.

## Introduction

Cervical cancer remains the fourth most common cancer in women worldwide, affecting 13.1 of every 100,000 women [1]. Cervical cancer incidence and mortality rates in Latin America and the Caribbean are around 30% to 40% higher than in non-Hispanic white women [2]. Hispanic women living in Puerto Rico have a higher incidence of cervicovaginal cancer (12.9 vs. 7.2 per 100,000), as well as a higher death rate when compared to women in the U.S. mainland (2.2 vs. 2.0 per 100,000) [2, 3]. A recent study confirmed that women living in Puerto Rico have the highest age-adjusted incidence of cervical cancer in the U.S., with an increasing incidence from 2001 to 2017 from 9.2 to 13 per 100,000 person-years [4].

Cervical carcinogenesis is a complex process influenced by the host and the microbiota [5]. One key element of cervical cancer development and an etiological agent of the disease is the persistence of human papillomavirus (HPV) [6, 7]. HPV, particularly high-risk HPV (HR-HPV) genotypes, increases the risk of cervical neoplasia [8]. These cervical abnormalities are detected through cytology, which provides a way to identify different squamous cell changes known as atypical squamous cells of undetermined significance (ASCUS), low-grade or high-grade squamous intraepithelial lesions (LGSIL and HGSIL, respectively) [9].

Recent evidence indicates that the cervicovaginal environment, including the microbiota, may play a role in viral persistence and the progression of epithelial lesions that lead to carcinogenesis [10]. Differential activation of the mucosal immune system by varying cervicovaginal microbiota composition may play a role in lesion development [11]. The composition of the cervicovaginal microbiota is influenced by ethnicity, age and lifestyle. For example, the cervicovaginal microbiota of White women is dominated by *Lactobacillus crispatus*, while Hispanic and Black women have a higher prevalence of non-*Lactobacillus* genera such as *Dialister, Gardnerella, Clostridium, or Prevotella* [12]. Women living in Puerto Rico (mainly Puerto Ricans) are Hispanics with a genetic admixture of European, Amerindian, and African ancestry. This population, however, differs from U.S. and Mexican Hispanics due to a greater contribution of African genetic ancestry (<6.2% vs. 16-32%) [13-15]. The cervicovaginal microbiota of women living in Puerto Rico reproductive-age women is dominated by *L. iners* [16]. In addition to ancestry, physiological, lifestyle, and hormonal changes throughout a woman’s life can significantly influence the vaginal microbiome [17]; however, the cervicovaginal microbiota of pregnant and menopausal Puerto Rican women has not yet been characterized.

We posited that the cervicovaginal microbial diversity of Hispanic women residing in Puerto Rico would vary with pregnancy, menopause, and HPV infection state. We found that *L. iners* and other diverse microbial cervicovaginal taxa were commonly found in Puerto Rican women regardless of their physiological stage, and that these microbes, in combination with HR-HPV may promote a favorable environment for cervical lesion development.

## Results

### Population description

A total of 333 samples from 294 women were included in the analyses (**Table S1**). This study cohort included three groups of women: reproductive-age non-pregnant (N=196), pregnant (N=37), and menopausal (N=61) that differed in terms of the severity of cervical lesions and HPV infection state (**Table 1**).

**Table 1.** Study population description by women’s group.

| Variables | Non pregnant<br>(N=196) | Pregnant<br>(N=37) | Menopause<br>(N=61) | TOTAL<br>(N=294) | P value & |
| --- | --- | --- | --- | --- | --- |
| **Age** |  |  |  |  | *<1e-16* |
| Mean (SD) | 36.0 (8.71) | 28.3 (4.69) | 52.1 (7.25) | 38.4 (11.0) |  |
| Median [Min, Max] | 35.5 [21.0, 54.0] | 28.0 [21.0, 43.0] | 54.0 [28.0, 60.0] | 37.0 [21.0, 60.0] |  |
| **BMI category** |  |  |  |  | *0.003* |
| Normal | 79 (40.3%) | 7 (18.9%) | 18 (29.5%) | 104 (35.4%) |  |
| Underweight | 4 (2.0%) | 1 (2.7%) | 0 (0%) | 5 (1.7%) |  |
| Overweight | 39 (19.9%) | 12 (32.4%) | 26 (42.6%) | 77 (26.2%) |  |
| Obese | 73 (37.2%) | 17 (45.9%) | 16 (26.2%) | 106 (36.1%) |  |
| Missing | 1 (0.5%) | 0 (0%) | 1 (1.6%) | 2 (0.7%) |  |
| **Vaginal pH** |  |  |  |  | *1e-8* |
| Mean (SD) | 5.40 (0.540) | 5.05 (0.522) | 5.67 (0.469) | 5.42 (0.549) |  |
| **Cervical lesions &&** | 67 (34.2%) | 20 (54.1%) | 13 (21.3%) | 100 (34.0%) | *0.001* |
| **Type of cervical lesion &&&** |  |  |  |  | *0.007* |
| HGSIL | 33 (16.8%) | 12 (32.4%) | 6 (9.8%) | 51 (17.3%) |  |
| LGSIL | 34 (17.3%) | 8 (21.6%) | 7 (11.5%) | 49 (16.7%) |  |
| NILM | 114 (58.2%) | 13 (35.1%) | 45 (73.8%) | 172 (58.5%) |  |
| Missing | 15 (7.7%) | 4 (10.8%) | 3 (4.9%) | 22 (7.5%) |  |
| HPV Status | 151 (77.0%) | 27 (73.0%) | 42 (68.9%) | 220 (74.8%) | *0.444* |
| **HPV type &&&&** |  |  |  |  | *0.300* |
| Only HR-HPV | 84 (42.9%) | 21 (56.8%) | 28 (45.9%) | 133 (45.2%) |  |
| Only LR-HPV | 17 (8.7%) | 1 (2.7%) | 2 (3.3%) | 20 (6.8%) |  |
| Both | 49 (25.0%) | 5 (13.5%) | 11 (18.0%) | 65 (22.1%) |  |
| Negative | 44 (22.4%) | 10 (27.0%) | 18 (29.5%) | 72 (24.5%) |  |
| Missing | 2 (1.0%) | 0 (0%) | 2 (3.3%) | 4 (1.4%) |  |
| **HPV type** |  |  |  |  | *0.441* |
| HR-HPV type | 133 (67.9%) | 26 (70.3%) | 39 (63.9%) | 198 (67.3%) |  |
| Only LR-HPV types | 17 (8.7%) | 1 (2.7%) | 2 (3.3%) | 20 (6.8%) |  |
| Negative | 44 (22.4%) | 10 (27.0%) | 18 (29.5%) | 72 (24.5%) |  |
| Missing | 2 (1.0%) | 0 (0%) | 2 (3.3%) | 4 (1.4%) |  |
| **Number of HR-HPV genotypes** |  |  |  |  | *0.654* |
| Mean (SD) | 1.77 (0.953) | 1.54 (0.706) | 1.95 (1.39) | 1.77 (1.03) |  |
| **Number of LR-HPV genotypes** |  |  |  |  | *0.159* |
| Mean (SD) | 1.24 (0.432) | 1.00 (0) | 1.07 (0.267) | 1.20 (0.401) |  |
| **HPV Vaccine** | 40 (20.4%) | 5 (13.5%) | 3 (4.9%) | 48 (16.3%) | *0.006* |
| **Alcohol drinking** | 99 (50.5%) | 1 (2.7%) | 28 (45.9%) | 128 (43.5%) | *1e-8* |
| **Tabaco smoking** | 27 (13.8%) | 4 (10.8%) | 4 (6.6%) | 35 (11.9%) | *0.311* |
| **Heterosexual preference** | 191 (97.4%) | 36 (97.3%) | 61 (100%) | 288 (98.0%) | *0.561* |
| **Number of sexual partners** |  |  |  |  | *1e-8* |
| Mean (SD) | 4.94 (4.15) | 4.82 (4.72) | 2.36 (1.39) | 4.38 (3.95) |  |
| **Antibiotic intake last 2 months** | 12 (6.1%) | 3 (8.1%) | 6 (9.8%) | 21 (7.1%) | *0.549* |
| **Pregnancy trimester** |  |  |  |  | *1.000* |
| 1rst | - | 8 (21.6%) | - | 8 (2.7%) |  |
| 2nd | - | 20 (54.1%) | - | 20 (6.8%) |  |
| 3rd | - | 9 (24.3%) | - | 9 (3.1%) |  |

& Fisher's Exact Test and Kruskal-Wallis test for categorical and continous variables respectively
&& Positive samples includes the HGSIL or LGSIL
&&& Samples diagnosed as ASCUS and HPV positive were consider LGSIL
&&&& HR-HPV: high-risk HPV, LR-HPV: low-risk HPV

We first compared the demographic and health characteristics of women in these three groups. The median age was 35.5 for non-pregnant, 28 for pregnant and 54 for menopause. The Body Mass Index (BMI) for the non-pregnant women was mostly normal (40.3% of women), while most menopausal women were overweight (42.6%, P<0.003). Pregnant women showed the highest prevalence of High grade squamous intraepithelial lesion (HGSIL; 32.4%), while menopausal women had the lowest (9.8%). HPV infections were present in most subjects (74.8%) with no differences among groups of women. Only 16.3% of women were HPV vaccinated, mainly in the non-pregnant group (20.4%). Regular alcohol drinking occurred in 43.7% of the women, mostly in non-pregnant and menopausal women, and only one pregnant woman reported drinking regularly (2.7%). Tabaco smoking was less common (11.9%) and similarly distributed among groups. Most subjects were self-declared heterosexuals (98.0%) and reported a sexual history of 3 sexual partners, which was higher in non-pregnant women (median: 4 sexual partners). The use of antibiotics for the last two months was reported only in 7.1% of women, and 1.0% did not answer. These women were retained in the study since they were uniformly distributed among women’s groups. However, this variable was always included in the models as covariable. Pregnant women were mostly in their 2^nd^ pregnancy trimester (54.1%), followed by 3^rd^ (24.3%) and 1^st^ (21.6%, **Table 1**).

### Cohort description according to cervical lesion and HPV status

We compared the prevalence of HPV infection and cervical lesions across women’s groups. We found an overall prevalence for cervical lesions of 34.0% (100/294) (**Table 1**), with 17.3% of HGSIL and 16.7% of Low grade squamous intraepithelial lesion (LGSIL). For any HPV genotype infection (HPV status), we found an overall prevalence of 74.8% (220/294), where 67.3% (198/294) of patients were HR-HPV positive. We also reported the prevalence of infections with only HR-HPV (45.2%, 133/294), with only LR-HPV (6.8%, 20/294) and mixed infections (both HR and LR-HPV types at the same time, 22.1%, 65/294, **Table 1**). Mixed infections were higher in LGSIL (30.6%) than in HGSIL (9.8%, P=0.031, data not shown). No differences in HGSIL prevalence were observed comparing patients with single HPV 16 infection than when HPV 16 was accompanied by another HPV genotype (P>0.050). As expected, HR-HPV prevalence was higher in HGSIL (76.5%) compared to in LGSIL and negative cervical lesions (*Padj*<1e-4, **Table 2**). HGSIL samples showed mostly infections with HR-HPV 16 (37.8%), HPV-HR 52 (28.9%) and HPV-HR 51 genotypes (24.4%) (**Table 3**). Interestingly, no HGSIL and only one LGSIL sample were related to HR-HPV-18 infections (**Table 3**). In terms of the number of HPV types, women had a median of 1 (mean of 1.8, max: 6) HR-HPV genotype and 1 (mean of 1.2, max: 2) LR-HPV genotype, with no difference among women groups (**Table 1**) or between lesion types (HR-HPV, *P*=0.195; LR-HPV, *P*=0.857, Kruskal-Wallis test, **Table 2**). The most prevalent HR-HPV genotypes were 51 (31.4%), 16 (20.9%), 33 (16.8%), 52 (14.5%) and 56 (14.5%), with similar prevalence among women’s groups (*P>0*.*246*). Only HR-HPVs 59 and 31 were marginally more prevalent in menopausal women (11.9 and 9.5%, respectively, *P<0*.*024, Padj>0*.*173*, **Table S2**). Among the LR-HPV genotypes 53 (22.7%), 74 (10.0%), and 44 (4.1%) were the most prevalent and not different among women groups.

**Table 2.**
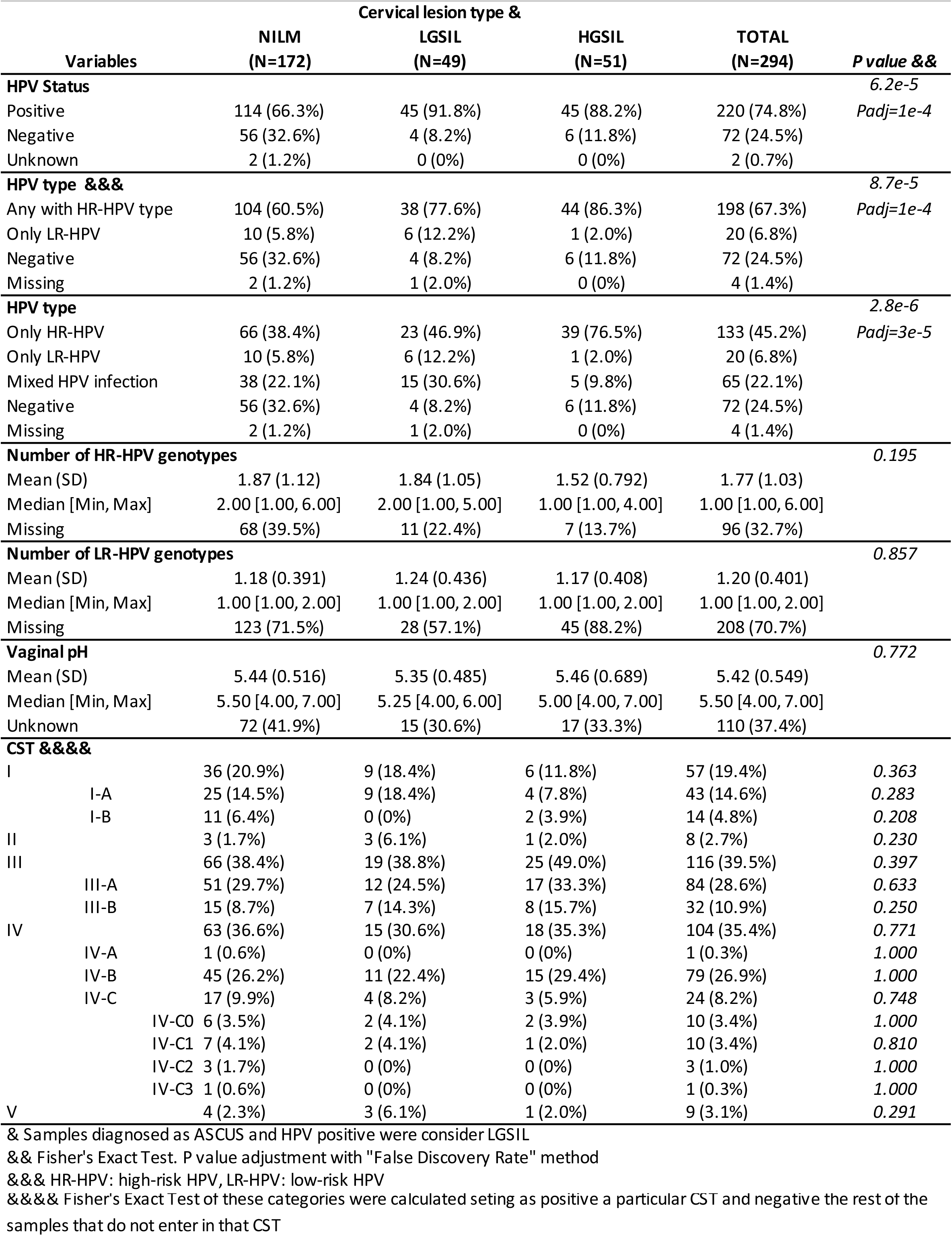
HPV infection prevalence, HPV number of genotypes and Community State Types (CSTs) among women with differential cervical lesion statuses

**Table 3.** Prevalence of HPV genotypes per cervical lesion groups including all women.

| HPV types | HGSIL<br>(N=45) | LGSIL<br>(N=45) | NILM<br>(N=114) | TOTAL<br>(N=220) | *P value &* |
| --- | --- | --- | --- | --- | --- |
| HR51 | 11 (24.4%) | 13 (28.9%) | 42 (36.8%) | 69 (31.4%) | 0.301 |
| HR16 | 17 (37.8%) | 7 (15.6%) | 19 (16.7%) | 46 (20.9%) | 0.012 (Padj=0.092) |
| HR33 | 6 (13.3%) | 7 (15.6%) | 23 (20.2%) | 37 (16.8%) | 0.580 |
| HR52 | 13 (28.9%) | 3 (6.7%) | 14 (12.3%) | 32 (14.5%) | 0.009 (Padj=0.092) |
| HR56 | 4 (8.9%) | 10 (22.2%) | 16 (14.0%) | 32 (14.5%) | 0.193 |
| HR66 | 0 (0%) | 5 (11.1%) | 19 (16.7%) | 29 (13.2%) | 0.005 (Padj=0.092) |
| HR45 | 5 (11.1%) | 3 (6.7%) | 7 (6.1%) | 16 (7.3%) | 0.580 |
| HR35 | 5 (11.1%) | 2 (4.4%) | 7 (6.1%) | 15 (6.8%) | 0.519 |
| HR68 | 0 (0%) | 5 (11.1%) | 10 (8.8%) | 15 (6.8%) | 0.055 |
| HR39 | 1 (2.2%) | 2 (4.4%) | 8 (7.0%) | 12 (5.5%) | 0.593 |
| HR58 | 2 (4.4%) | 4 (8.9%) | 4 (3.5%) | 11 (5.0%) | 0.408 |
| HR59 | 0 (0%) | 0 (0%) | 8 (7.0%) | 8 (3.6%) | 0.044 (Padj=0.202) |
| HR31 | 0 (0%) | 2 (4.4%) | 3 (2.6%) | 5 (2.3%) | 0.407 |
| HR18 | 0 (0%) | 1 (2.2%) | 1 (0.9%) | 2 (0.9%) | 0.689 |
| LR53 | 4 (8.9%) | 12 (26.7%) | 32 (28.1%) | 50 (22.7%) | 0.023 (Padj=0.132) |
| LR74 | 3 (6.7%) | 6 (13.3%) | 11 (9.6%) | 22 (10.0%) | 0.592 |
| LR44 | 0 (0%) | 4 (8.9%) | 4 (3.5%) | 9 (4.1%) | 0.105 |
| LR6 | 0 (0%) | 1 (2.2%) | 2 (1.8%) | 8 (3.6%) | 1.000 |
| LR70 | 0 (0%) | 1 (2.2%) | 4 (3.5%) | 6 (2.7%) | 0.826 |
| LR54 | 0 (0%) | 1 (2.2%) | 2 (1.8%) | 4 (1.8%) | 1.000 |
| LR40 | 0 (0%) | 1 (2.2%) | 1 (0.9%) | 2 (0.9%) | 0.689 |
| LR19 | 0 (0%) | 0 (0%) | 1 (0.9%) | 1 (0.5%) | 1.000 |
| LR43 | 0 (0%) | 0 (0%) | 1 (0.9%) | 1 (0.5%) | 1.000 |

& Fisher's Exact Test among total prevalence per cervical lesion group. P value adjustment with "False Discovery Rate" method
HR=high-risk HPV types, LR=low-risk HPV types

The cervicovaginal pH of non-pregnant women had a median of 5.5, which was significantly lower than in menopausal women (median pH 6.0, *Padj*<0.004, pairwise Wilcox-Test) but significantly higher than in pregnant women (median pH=5.0, *Padj*<0.006, pairwise Wilcox-Test, **Table 1, Fig. S1-a**). No significant difference in cervicovaginal pH was observed among cervical lesions (*P*=0.772, Kruskal-Wallis test, **Table 2**).

### Microbial Community Composition

The differences in composition and structure of cervicovaginal microbial communities were evaluated using PERMANOVA models. To avoid collinearity between cervicovaginal pH and women’s group, cervicovaginal pH was only included in models when stratified by women groups. We found that cervicovaginal microbiota composition significantly differed among women’s group (significant for unweighted, P=0.003; generalized, P=0.007; and weighted UniFrac, *P*<0.030; R^2^=0.029-0.032). We observed that microbiota from menopause differed from pregnant (*Padj*<0.014) and from non-pregnant subjects (*Padj*<0.038, for unweighted, and weighted UniFrac, **Fig. 1a**. and generalized UniFrac, not shown). The presence of cervical lesions or HPV infections did not significantly influence microbial composition among all women *(P*>0.050, R^2^<0.01). For the stratified analysis among non-pregnant or menopausal subjects, cervicovaginal pH was significantly different (P<0.01 among non-pregnant, for all metrics; P=0.046 only for unweighted UniFrac). Finally, for pregnant women, the microbial composition also differed by pregnancy trimester (for all distances, *P*<0.034, R^2^>0.13), where 1^st^ and 2^nd^ trimesters significantly differed (*P*<0.007, R^2^=0.15, pairwise PERMANOVA, for all distances, **Fig. 2-a, b**). We also did a higher rarefaction of 5,000 reads per sample and found similar results (**Fig. S2 a-e**). To provide a comprehensive analysis, we included data analyzed using both SILVA 138 and the Greengenes extended database. A comparison of the results is presented in **Fig. S6** and **S7**, with similar outcomes observed with both databases. However, we found no significant reduction of alpha diversity with the pregnancy trimesters when using SILVA compared to the significant result using GreenGenes. Apart from this, all diversity metrics were consistent between the two databases. In addition, we observed that Greengenes extended database proved to be better for resolving Lactobacillus species in the cervicovaginal environment.

**Fig 1.**
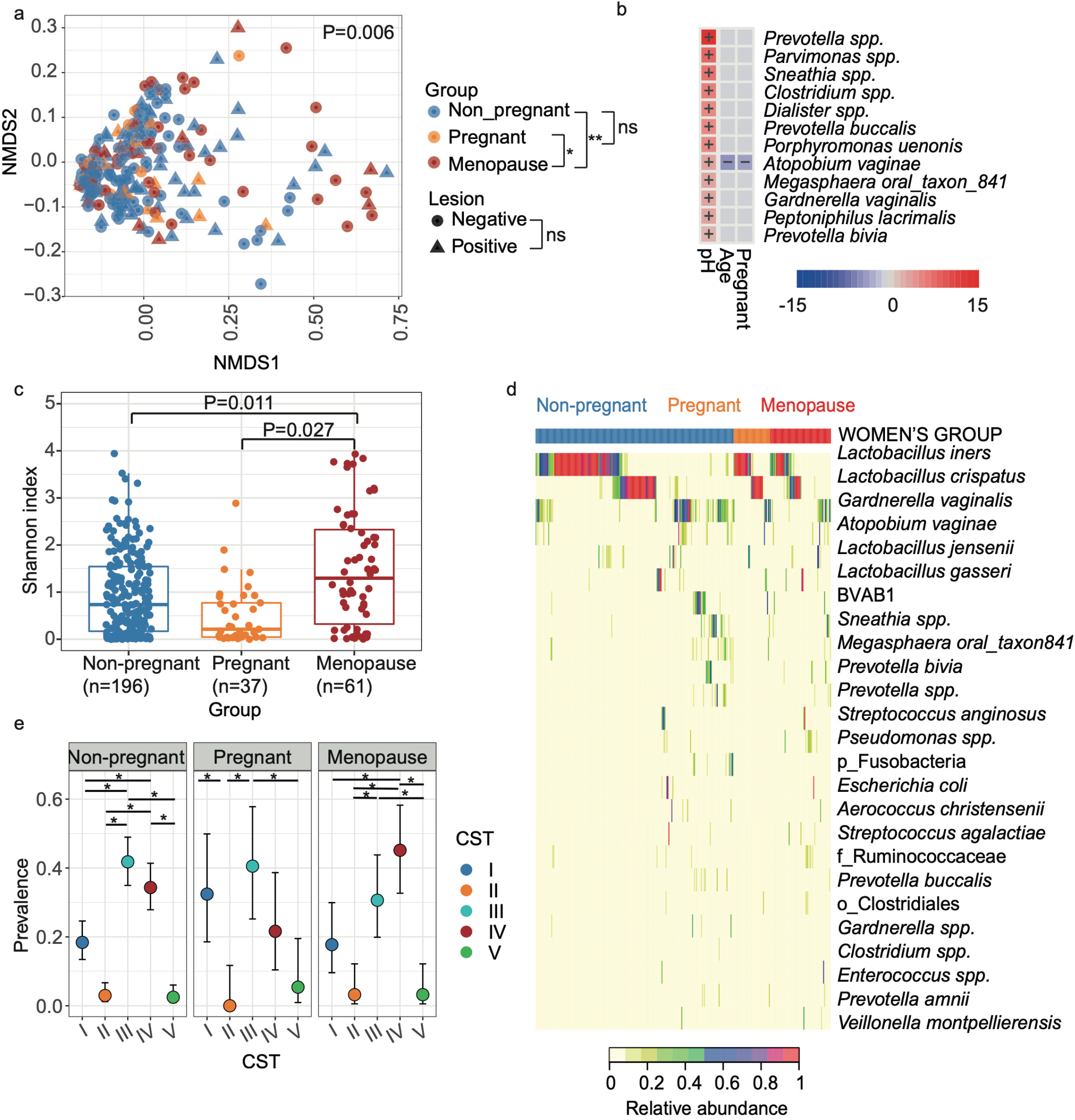
The cervicovaginal microbiota at the species level in non-pregnant, pregnant, and menopausal women were compared. (**a**) Beta diversity with unweighted UniFrac distance. Analysis was performed with PERMANOVA. (**b**) The heatmap summarizes all the significant relationships between microbial taxa and sample metadata. Color key: -log (q-value) * sign color key (coefficient). Cells indicating significant associations are colored (red or blue) and overlaid with a plus (+) or minus (-) sign indicating the direction of the association. **(c)** Shannon diversity among women groups, analysis was performed with Linear model. (**d**) Heatmap showing the 25 most abundant taxa, ordered by women’s group and hierarchical clustering within women’s groups. (**e**) Prevalence of Community State genotypes (CSTs) by women’s group. Analysis was performed with fisher’s exact test (*P<0.05, **P<0.005). Panels created in *R* (see methods); and final multi-panel figure mounted using Illustrator Adobe Inc. (2021); https://adobe.com/products/illustrator.

**Fig. 2.**
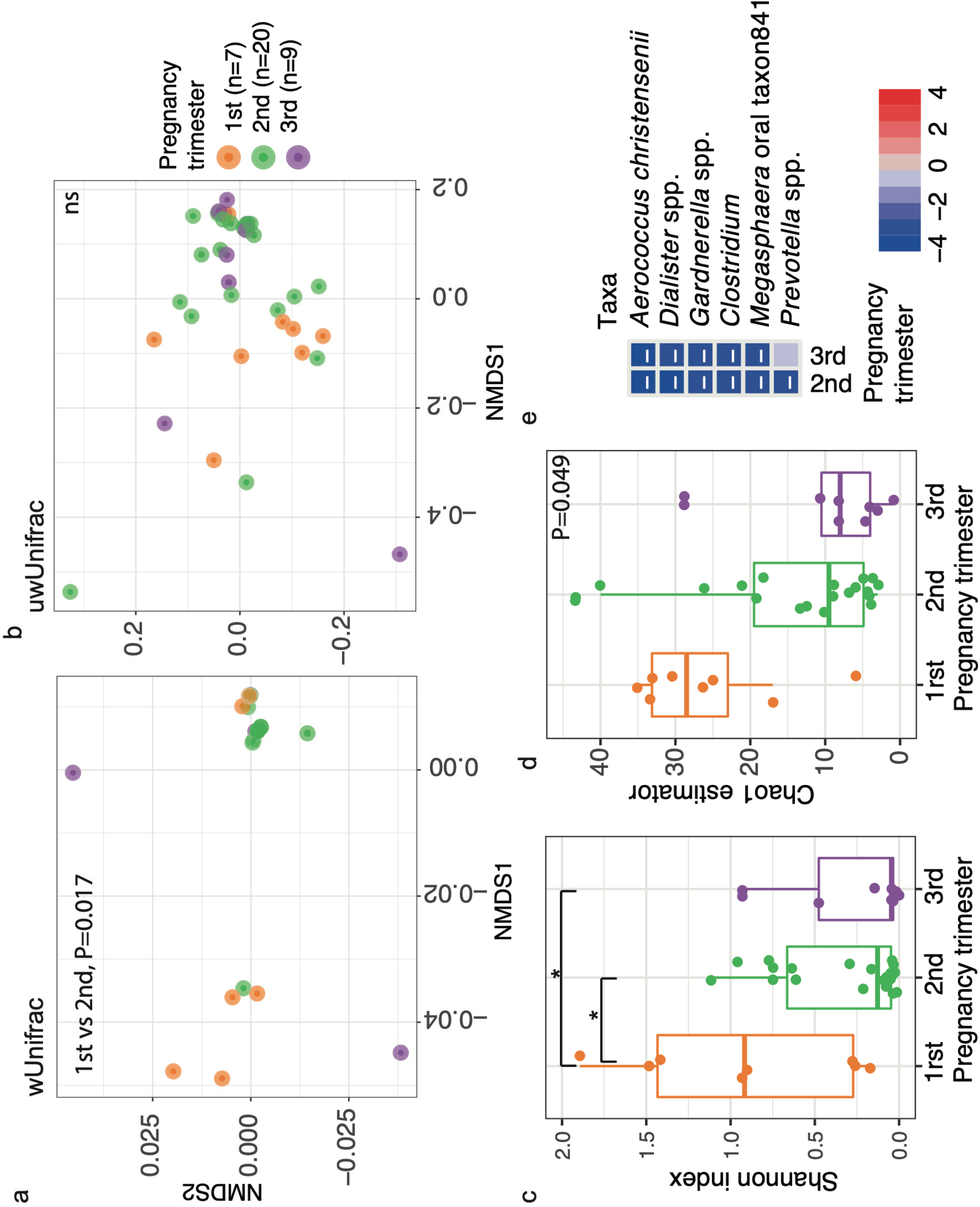
Cervicovaginal microbiota at the species level changes with pregnancy trimesters. (**a, b**) Beta diversity for (**a**) weighted and (**b**) unweighted Unifrac distances. Analysis was performed with PERMANOVA. (**c, d**) Alpha diversity for Shannon index (**c**) and Chao1 estimator (**d**). Analysis was performed with Linear model. (**e**) Heatmap of taxa significantly associated with pregnancy trimester analyzed with Maaslin2. The heatmap summarizes all the significant relationships between microbial taxa and sample metadata. Color key: -log (q-value) * sign color key (coefficient). Cells indicating significant associations are colored (red or blue) and overlaid with a plus (+) or minus (-) sign indicating the direction of the association, being the first trimester the reference. *P<0.05, ns=not significant. Panels created in *R* (see methods); and final multi-panel figure mounted using Illustrator Adobe Inc. (2021); https://adobe.com/products/illustrator.

### Association of bacterial taxa with women’s group, cervicovaginal pH, HPV infections, and cervical lesions

We also analyzed the taxa association, including in the model the covariables of interest using Maaslin2 algorithm, which relies on linear models. We found that overall, *Prevotella, Parvimonas* spp., *Sneathia, Clostridium, Dialister, Prevotella buccalis, Porphyromonas uenonis, Megasphaera oral taxon841, Peptoniphilus lacrimalis, Gardnerella vaginalis, Atopobium vaginae*, and *Prevotella bivia* had significantly greater proportions in less acidic cervicovaginal pH (*Padj*<0.050, **Fig. 1b, Table S3**). Also, being pregnant or younger was associated with lower *Atopobium vaginae* (**Fig. 1b, Table S3**). A separate analysis was performed for pregnant women by pregnancy trimester and observed that *Aerococcus christensenii, Dialister spp*., *Gardnerella spp*., *Clostridium, Megasphaera oral taxon841* decreased in proportion consistently in 2^nd^ and 3^rd^ when compared to the 1^st^ trimester, and *Prevotella* spp. decreased in the 2^nd^ trimester (*Padj*>0.046, **Fig. 2e, Fig S2 i**). In general, anaerobic opportunistic taxa were positively associated with higher pH, and negatively associated with age and pregnancy.

We did not detect any bacterial taxa significantly associated with HPV infection or cervical lesions (*Padj*>0.104, Maaslin2). However, before *P-value* adjustment, two taxa had increased proportion in low-risk HPV infections compared to negative samples: *Corynebacterium* spp. and *Methanobrevibacter* spp. (*P=*0.006, and *P*=0.030, respectively). Also, two taxa were associated with HR-HPV infections before the P value adjustment for multiple comparisons: *Ureoplasma* decreased while *Clostridium* spp. Increased in proportion (*P*=0.035, and *P*=0.043), and, for cervical lesions *Escherichia coli*, increased with LGSIL (*P=0*.*007*; Maaslin2). Although marginally, HPV infections and cervical lesions was associated with an increase of the relative abundance of anaerobic-opportunistic taxa, and the significance of these associations may increase with a larger cohort size.

### Alpha diversity varies by group and HGSIL

We used linear mixed models (LMM), including the same covariables of interest as we employed for the beta diversity and taxa association analyses. We found that women’s group (*P*<6.2e-5) and cervicovaginal pH (*P*<0.002) were significantly associated with alpha diversity (**Fig. S1-C**). Within women’s group, menopausal subjects showed greater Shannon diversity than non-pregnant (*Padj*<0.010, least-squares means (EMM) for Shannon and Chao1) and then pregnant women (*Padj*=0.027, only for Shannon, EMM) with no differences between non-pregnant and pregnant subjects (**Fig. 1-c**). However, no significant alpha diversity differences were observed between cervical lesion severity levels.

For the stratified analyses for each women’s group we found that for non-pregnant group, alpha diversity was greater at higher cervicovaginal pH or age, and in the presence of cervical lesions (HGSIL or LGSIL) compared with healthy subjects (*P*<0.050, pairwise *Padj*<0.050, for Shannon, **Fig. S1-b**, and Chao1, P=0.004, Padj<0.003, EMM). The later comparisons within pregnant (Padj>0.390, **Fig. S1c**) or menopause women groups were not different (P>0.450, **Fig. S1d**). Instead, HPV infection did not significantly influence alpha diversity among all women or non-pregnant women (*P*>0.050, LMM).

A similar analysis was performed per trimester of pregnancy, initially showing no significant association (*P*>0.050, for Shannon and Chao1). Nonetheless, when the two outliers were top-coded (data points > 2x standard deviation away from the mean within each trimester, were assigned the last highest value), the pregnancy trimester variable showed a significant tendency to decrease late in pregnancy (*P*<0.049, for both alpha metrics). Pairwise analysis showed that 1^st^ trimester was significantly greater than the 2^nd^ and 3^rd^ trimester (*P*<0.010, for Shannon diversity, not significant for Chao1 estimator, **Fig. 2c, d**). Indeed, when considering a 5,000 read rarefaction level we found that only the Chao1 estimator evidenced a significant decrease in richness from the first to the third pregnancy trimesters (*P*=0.01, **Fig S2 f, g**).

In general, alpha diversity analyses showed the greatest values in women with cervical lesions or menopause, and was lowest in pregnant women, particularly in the last trimester of pregnancy.

### Women living in Puerto Rico are dominated by diverse microbial Community State Genotypes

We also evaluated the Community State genotypes (CST) distribution among women’s groups. CSTs are categories that allow classifying each woman’s cervicovaginal microbial composition. In general, they are defined by the dominance of *L. crispatus* (CST-I), *L. gasseri* (CST-II), *L. iners* (CST-III), *L. jensenii* (CST-V), or non-*Lactobacillus* dominated (CST-IV) [12]. A finer classification by CST sub-groups has also been proposed and used in this analysis [18]. We used fisher’s exact test to test the prevalence of these CSTs among women’s groups. As a side note, when analyzing the sequences, only a small percentage (0.61%) of the *Lactobacillus* ASV’s were unable to be identified at the species level, indicating that our data for this analysis is reliable. We observed that most women had *Lactobacillus*-dominated CSTs (64.9%), prevailing CST-III (39.5%). No significant differences were observed among women’s groups (*P*=0.125). However, when stratifying by women’s group, non-pregnant women had a higher prevalence of CST-III (42.3%) followed by CST-IV (34.7%) than any other CST (*Padj*<0.010, **Fig. 1-e**). Instead, pregnant women showed higher CST-III (40.5%) and CST-I (32.4%, *Padj*<0.010, **Fig. 1-e**), with a higher prevalence of the protective *L. crispatus* (**Fig. 1-d**). Non-pregnant and menopause women had mostly *L. iners* and CSTIII and CST-IV (**Fig. 1-d**). Indeed, for menopausal women, CST-IV (45.9%) followed by CST-III (29.5%) were the most prevalent (*Padj*<0.010, **Fig. 1-e**). Among the CST-IV, the subtype CST IV-C was strongly higher in menopausal women (24.6%) than in the other women’s groups (<4.1%, *Padj*=0.002, **Table 4**). This profile is characterized by a low relative abundance of *Lactobacillus spp*., *G. vaginalis, A. vaginae*, and BVAB1. Instead, a diverse array of facultative and strictly anaerobic bacteria exists. Among the different CST IV-C subtypes, menopausal women showed a higher prevalence of CST IV-C0 (11.5%) which is an even microbial profile with a moderate amount of *Prevotella*, and CST IV-C1 (9.8%) which is a *Streptococcus*-dominated profile (**Table 4, Fig. S1-d**).

**Table 4.** Distribution of CST categories among women’s group.

| CST categories | Women's group, n (%) |  |  |  | P value & |
| --- | --- | --- | --- | --- | --- |
|  | Non pregnant<br>(N=196) | Pregnant<br>(N=37) | Menopause<br>(N=61) | TOTAL<br>(N=294) |  |
| **I** | 34 (17.3) | 12 (32.4) | 11 (18.0) | 57 (19.4) | 0.161 |
| **I-A** | 27 (13.8) | 10 (27.0) | 6 (9.8) | 43 (14.6) | 0.071 |
| **I-B** | 7 (3.6) | 2 (5.4) | 5 (8.2) | 14 (4.8) | 0.449 |
| **II** | 6 (3.1) | 0 (0) | 2 (3.3) | 8 (2.7) | 0.746 |
| **III** | 83 (42.3) | 15 (40.5) | 18 (29.5) | 116 (39.5) | 0.301 |
| **III-A** | 59 (30.1) | 15 (40.5) | 10 (16.4) | 84 (28.6) | 0.021 Padj=0.084 |
| **III-B** | 24 (12.2) | 0 (0) | 8 (13.1) | 32 (10.9) | 0.034 Padj=0.109 |
| **IV** | 68 (34.7) | 8 (21.6) | 28 (45.9) | 104 (35.4) | 0.057 |
| **IV-A** | 1 (0.5) | 0 (0) | 0 (0) | 1 (0.3) | 1.000 |
| **IV-B** | 59 (30.1) | 7 (18.9) | 13 (21.3) | 79 (26.9) | 0.226 |
| **IV-C** | 8 (4.1) | 1 (2.7) | 15 (24.6) | 24 (8.2) | 1e-7 Padj=2e-4 |
| **IV-C0** | 2 (1.0) | 1 (2.7) | 7 (11.5) | 10 (3.4) | 0.001 Padj=0.009 |
| **IV-C1** | 4 (2.0) | 0 (0) | 6 (9.8) | 10 (3.4) | 0.014 Padj=0.075 |
| **IV-C2** | 2 (1.0) | 0 (0) | 1 (1.6) | 3 (1.0) | 0.700 |
| **IV-C3** | 0 (0) | 0 (0) | 1 (1.6) | 1 (0.3) | 0.330 |
| **V** | 5 (2.6) | 2 (5.4) | 2 (3.3) | 9 (3.1) | 0.448 |

& Fisher's Exact Test. P value adjustment with "False Discovery Rate" method. Analyses were performed by row, setting in the contingency table positive and negative cases for a particular CST, being the negative cases all subjects that were not assigned to the evaluated category

In terms of cervical lesions (presence and types), analyses performed with all women by CST had no significant association (*P*<0.393, fisher’s exact test, **Table 2**), nor when stratifying by pregnant, non-pregnant or menopausal women (**Fig. 3, Fig. S4**). However, CSTs prevalence were significantly different among cervical lesions within non-pregnant women (positive/negative, *P*=0.033, **Table S4**), where the presence of cervical lesion showed lower CST-I prevalence compared to its absence, although not significantly (8.7% vs 22.3%, fisher’s exact test, *Padj*>0.137, data not shown). Analyses of subCSTs within non-pregnant women suggest that women negative for lesion showed higher prevalence of *Lactobacillus*-dominant groups CST I-A (*L. crispatus*-dominated) and CST-III-A (*L. iners*-dominated), although not significantly (**Fig. S4**). Instead, women with HGSIL showed higher prevalence of CST IV-B (characterized by having a high to the moderate relative abundance of *G. vaginalis* with some *Atopobium vaginae*), also not significantly. This CST-IV is characterized by, markers of cervicovaginal dysbiosis (**Fig. S4**).

**Fig 3.**
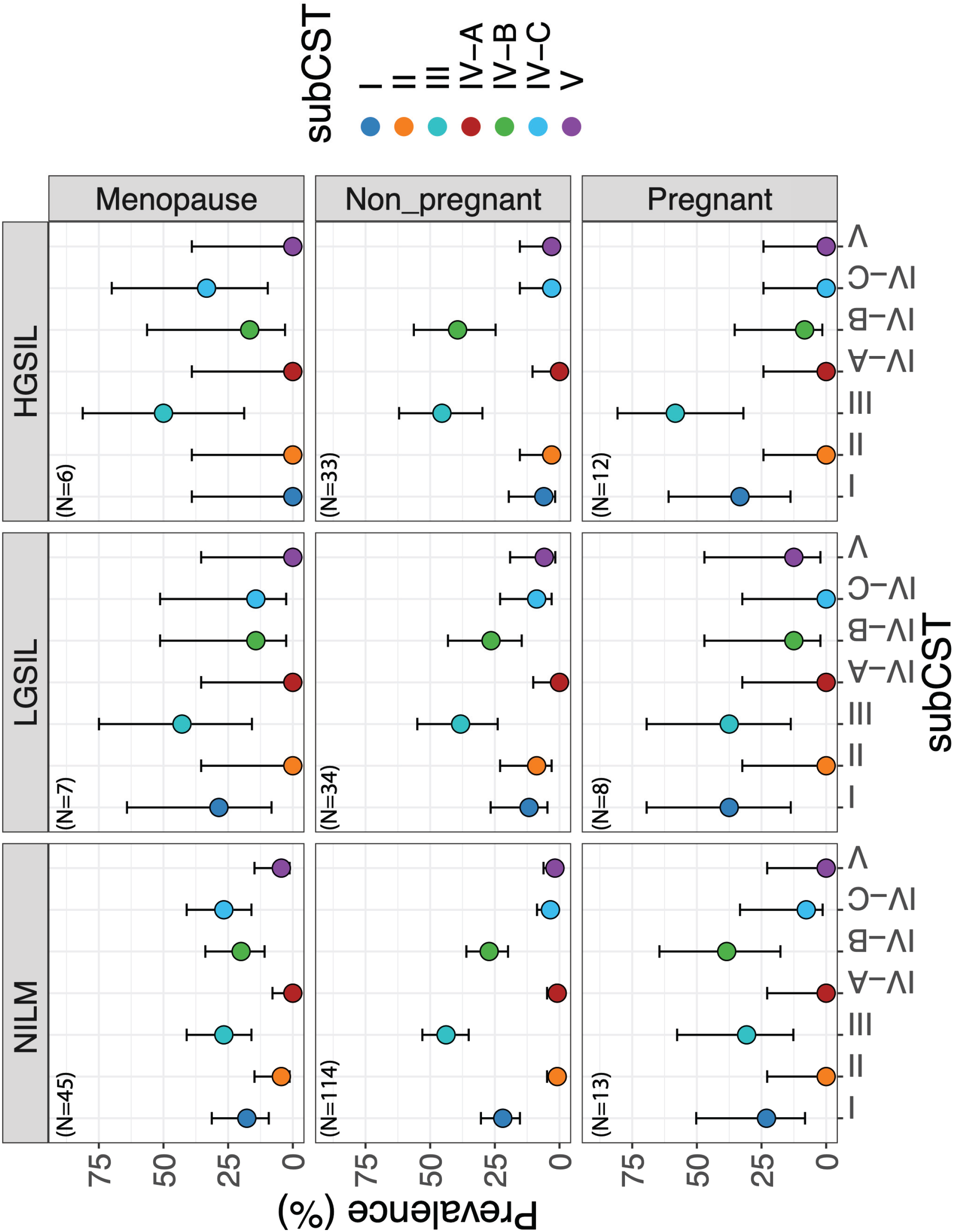
Community State Type (CST) distribution by cervical lesion genotypes among women groups. Bars correspond to 95% confidence intervals. CST prevalence was obtained by calculating the 100% among each women group and lesion type. Sample sizes are written in the left-hand upper corner of each box.

We found no significant differences (*P*<0.05, fisher’s exact test, data not shown) in CSTs with HPV infections (presence and types) among all women. However, when considering only non-pregnant women, we did find significant differences between HR-HPV/Only_LR-HPV/negative (*P*=0.039, fisher’s exact test, **Table S5**), which shows that women with LGSIL had a particularly low prevalence of CST-IV compared to HGSIL and negative samples (11.8% vs. 35.5% and 43% respectively, **Table S5**). In general, being menopausal, having cervical lesions, and being infected with exclusively HR-HPV infections are traits associated with diverse microbial profiles. We also aimed to compare CSTs from non-pregnant Puerto Rican women with those of U.S. Hispanic women (self-declared) reported by Ravel, et al. [12]. We found no significant differences between Hispanics from the U.S. vs. Puerto Rican Hispanics (*P*>0.132, fisher’s exact test, **Table S6**). E.g., for CST IV, 38.1% and 34.7% respectively (*P=*0.881).

### Most abundant *L. iners*-ASVs were not associated with cervical lesions or HPV infection

We also tested if the top15 most abundant ASV classified as *L. iners* were associated with cervical lesions or their types. We did not find significance for any *L. iners*-ASVs by cervical lesion or HPV infection, not even by the severity of lesion or HPV risk genotype (P>0.050, Fisher’s exact test, data not shown).

### Longitudinal changes of the cervicovaginal microbiota in a subset of women

To explore whether microbial profiles in this population vary over time we analyzed two-time points longitudinal data. Thirty-nine women (non-pregnant, n=29; pregnant, n=1; menopausal, n=9) who had had a second sampling between 4 to 16 months after their first visit were analyzed. Approximately half of women (51.3%, 20/39) kept the same CST between visits, and only 12.8% (5/39) changed from a *Lactobacillus*-dominated (all CST-III) to a diverse profile (IV). In comparison, 12.8% (5/39) changed from a diverse (CST-IV) to *Lactobacillus*-dominated profile (CST-I and II). A total of 35.9% (14/39) of women changed between different combinations of *Lactobacillus*-dominated CSTs (CST-I, II, and III), with a no particular tendency and with no differences among women’s groups (**Fig. S5, a, b**). Cervical lesion phenotype was maintained between the two visits for 56.4% (22/39) of women. Meanwhile, HPV status coincided in 67.6% (25/37) of women from the first to the second visit (**Fig. S5-c, d**). Changes in CST based on cervical lesions or HPV infection was also explored; however, sample sizes were too small to perform a reliable comparison. Transitions from *Lactobacillus*-dominated to non-*Lactobacillus*-dominated profiles or vise-versa did not show any significant trend for cervical lesions or HPV status (*P*<0.225, fisher’s exact test, **Table S7, S8**).

## Discussion

This study revealed not only a diverse and volatile cervicovaginal bacterial microbiota, but also a different distribution of oncogenic HPVs in this Puerto Rican cohort. The most frequent genotypes were HPV51, 16, 33, 52 and 56, mostly in HGSIL lesions, with few to no HPV18. This seems to indicate that both the quadrivalent HPV vaccine (4vHPV) (targeting HPV genotypes 6, 11, 16, and 18) and the 9-valent HPV vaccine may have limited effectivity as they lack coverage of HPV51 and 56, HPV genotypes that are prevalent in our cohort. The role of multiple HR-HPV infections, specifically HPV51 and 56 in the aggravation of cervical lesions has not been studied in Puerto Rico and deserves further attention. We found that women in menopause had the lowest vaccination prevalence (4.9%), and this is expected as vaccination in younger women has better promotion and adherence. In Puerto Rico, HPV vaccination started at time of HPV vaccine approval in USA back in 2006. The vaccination focusses at the time on the younger population, and many were vaccinated after being sexually active. Thus, the low rate of vaccination in this cohort.

The prevalence of HPV in this study was similarly high (74.8%; HR-HPV, 67.3%) to that reported in a previous study in the Puerto Rican population by our team (84%; HR-HPV, 79%) [16], and in Hispanic and Amerindian populations (77%; HR-HPV, 65.9%) [19]. The prevalence in HPV in Puerto Rico was higher than that observed in other Hispanic populations, including Costa Ricans (carcinogenic, 35%) [20], Europeans (∼20%) [21], Japanese (∼20%) [22] and Nigerian (carcinogenic, 40%) [23] using the same SPF10/LiPA25 HPV detection method. The high prevalence of HR-HPV genotypes in our cohort is however not generalizable to the population of Puerto Rico, as clinics cover general gynecology and colposcopy, that is, some patients are referred for colposcopy, hence the higher prevalence of HR-HPVs. In the past, a population-based study in PR, showed that the prevalence of HR-HPV genotypes was lower, however, they only included only four HPVs 6, 11, 16 and 18 and used other detection method [24].

Although Puerto Rico is a Caribbean country, it differs from other Afro-Caribbean women from Barbados, where the cervicovaginal microbial community is mostly dominated by non-*Lactobacillus* genera (72%), including *Prevotella* [25]. It is possible this is because of differences in African ancestry between Barbados and Puerto Rico but could also result from differences in lifestyle. The cervicovaginal microbiota of women living in Puerto Rico were dominated by *L. iners* (39.5%) or diverse non-*Lactobacillus* communities (35.4%). This is consistent with previous work from our lab [16], U.S. Hispanics [12], and Venezuelans [26].

The high prevalence of *L. iners* and diverse profiles across all samples reflects a cervical microbial environment with low stability, that is, a dynamic ecosystem with frequent changes, particularly after perturbations, and therefore with a higher risk of developing adverse health outcomes [27, 28]. Transitions between *L. iners* and high-diversity microbial environments are the most frequent [29, 30]. Although *L. iners* is a *Lactobacillus* species, its metabolic and ecological characteristics make it the least protective cervicovaginal *Lactobacillus*. Unlike other health-associated *Lactobacillus, L. iners* can cohabit in a diverse environment and act as a vaginal symbiont, or as an opportunistic pathogen if surrounded by other anaerobic pathogens [30]. Its H _2_O _2_ production is null or very limited [31, 32]. It also does not promote a sufficiently acidic environment that inhibits undesirable bacteria [33]. Additionally, it generates only L-lactic acid instead of the protective D-lactic acid isomer. This promotes inflammation [34], and facilitates invasion of the cervix by opportunistic bacteria [35]. Diverse cervical microbial profiles have also been associated with high levels of pro-inflammatory genital cytokines. This damages the endocervix’s columnar epithelial barrier, exposing it to the colonization of other microbial or pathogenic agents [36, 37]. The normal healthy cervicovaginal microbial profile of women living in Puerto Rico without cervical lesions or HPV infections is already diverse [16], and are therefore more prone to cervicovaginal dysregulation and inflammatory conditions [38].

Diverse microbial profiles in our non-pregnant population were mostly represented by CST-IV-B (30.1%), like Hispanic women in the United States [18]. This profile is considered of higher risk for bacterial vaginosis since it includes taxa such as *Gardnerella vaginalis* [39] that are associated with an increased risk of viral infections including HPV. [40-44]. Menopausal women, besides having this profile (21.3%), also had a high prevalence of CST-IV-C (24.6%), with some women being high in *Prevotella* (CST-IV-C0) and others in *Streptococcus* (CST-IV-C1) as previously seen in other menopause studies in U.S. population [18, 45]. Although CST-IV-C is a non-lactobacillus-dominated profile, it has been associated with the lowest Nugent scores (Gram stain scoring system to diagnose bacterial vaginosis, where lower values suggest no B.V.). However, CST-IV-C high in *Prevotella* has been shown to induce a significantly greater pro-inflammatory signaling [36], and is considered unfavorable for menopausal women, particularly those with cervical lesions or infected by HPV. During menopause, significant hormonal changes occur. The decline in estrogen level can lead to a decrease in the cervical mucus (vaginal dryness) and glycogen deposition, the primary source of *Lactobacillus* [46]. Low *Lactobacillus* induces high cervicovaginal pH leading to an environment more susceptible to infections [47].

Contrary to menopause, pregnant women showed a decrease in microbial alpha diversity (particularly later in pregnancy) and an increase in *Lactobacillus*-dominated profiles (CST-III and CST-I), a phenomenon already reported by other groups [48]. This conversion of high to low alpha diversity is accentuated in pregnant Hispanic and black populations [49], two racial groups with high cervicovaginal microbial diversity. Variation in cervicovaginal community composition during pregnancy may also correspond with hormonal changes. The increase in estrogen levels promotes higher glycogen deposition in cervicovaginal epithelial cells [50]. *Lactobacillus* spp. utilize glycogen as the primary carbohydrate source to produce lactic acid, contributing to the protective effect of a low cervicovaginal pH (lysing bacteria other than *Lactobacillus*). Additionally, the dominance of *Lactobacillus* produces significant amounts of bacteriocins, contributing to colonization resistance [11, 51-53]. This defensive strategy explains the reduction of *Gardnerella* spp., *Atopobium vaginae* and *Prevotella* spp. observed in our study and reported in other pregnant Hispanics [49]; identifying a reduction in other BV-associated taxa we identified in our study, such as *Aerococcus christensenii, Megasphaera spp*., *Dialister spp*., and *Clostridium*. The lower abundance of *Atopobium vaginae* observed in pregnant women, as compared to women in menopause or the non-pregnant group, can be associated with the lower pH found during pregnancy which reduces the colonization of this BV-associated species. To note, in our pregnant cohort no women had the CST-III-B type, only CST-III-A (40.5%). CST-III-A is associated with a lower Nugent score and cervicovaginal pH [18], and, therefore greater protection against infection.

Previous studies have shown that greater microbial diversity in cervicovaginal samples and an *L. iners-*dominated community profile are both associated with an increased risk of HPV infection and persistence [45, 54-56]. However, in our study, very little or no association was detected. Differences with other studies may be explained by ethnicity or HPV detection method. However, this lack of a clear HPV-microbial association has been reported in other Hispanic populations with similar profiles and using the same HPV detection method [16, 26]. This might be due to the high HPV prevalence and a higher dominance of diverse and *L. iners*-dominated profiles in our cohort, when compared to Caucasian women with *L. crispatus-* dominated profiles. Additionally, differences between transient and persistent HPV infection may influence this association since only persistent infections have been associated with immune response and cervicovaginal dysbiosis [57]. Chronic cervical inflammation caused particularly by diverse bacterial populations, in addition to HPV persistence, creates a favorable context for neoplasia and later cancer development [58, 59]. However, longitudinal methods to define HPV persistence, especially using HPV typing methods with higher analytical sensitivity, are expensive and time-consuming, and therefore studies are scarce, but very relevant.

We found greater Shannon diversity in HGSIL and LGSIL and a lower prevalence of *L. crispatus*-dominated profiles (higher of CST III, IV-B and IV-C) when compared to healthy subjects, which supports previous findings [16, 60, 61]. Additionally, women with LGSIL tend to have coinfections with both HPV genotypes (HR and LR-HPV), while those with HGSIL showed a greater prevalence of infections with exclusively HR-HPV types. The coexistence of HR and LR-HPV risk genotypes were similarly prevalent in HR-HPV or LR-HPV exclusive infections, which has also been observed in Hispanic-Venezuelan populations [19]. This association has previously been linked to sexual partner turnover [62] and a lower risk of cervical cancer development [63-65].

We also explored the temporal stability between cervical lesions, HPV status, and cervicovaginal microbiota in a small subset of women. Changes observed between positive and negative lesions or HPV infections may be due to treatment (Loop Electrosurgical Excision Procedure, LEEP, or Endocervical curettage, ECC) or natural clearance. For the cervicovaginal microbiota, we found no specific trends between CST changes over time nor concerning HPV infection and cervical lesion. A limitation of this analysis is the long period between study visits for these patients. CST transition can be very rapid, for example switching for only a single day [66]. However, as in previous studies, CST-IV-B (with high to moderate *G. vaginalis* and *Atopobium vaginae*, present in 26.9% of the studied population) and CST-III, maybe more likely to transition to other CSTs [30, 66]. We found that similar unstable transition probabilities have already been reported by others, particularly from CST-III to CST-IV profiles in white and black population [30]. However, even in this small subset of women, we found that very few women maintain *Lactobacillus* dominated profiles, being more likely to progress to diverse profiles. Moreover, studies with symptomatic or asymptomatic BV subjects or women using cervicovaginal douches have demonstrated the transition from CST-III to dysbiosis-associated CST-IV-B is the most probable [29, 30]. The similarity among both profiles and the high prevalence of these in our study population, even in women without high-grade cervical lesions or oncogenic HPV types, highlights the volatile cervicovaginal microbial environment for Puerto Rican women, which may in part explains the high rates of cervical cancer in this population.

In conclusion, women living in Puerto Rico, regardless of their physiological stage, predominantly maintained cervicovaginal communities dominated by *L. iners* (CST III) or a diverse microbial profile (e.g., CSTs IV-B and IV-C). Even though these profiles have been previously associated with reduced cervicovaginal health, they were not found to be significantly linked to cervical lesion in our cohort. However, the high prevalence of HR-HPV and the presence of these less stable, non-optimal bacterial profiles may be associated with the greater risk of cervical cancer observed in our populations. As a result, Puerto Rican women may be considered a susceptible population for cervical lesion development.

## METHODS

### Study Design and participant sample collection

This cross-sectional study of adult women coming for gynecology and colposcopy evaluation at the UPR and San Juan City clinics (San Juan Puerto Rico, Metropolitan area), who did not meet the exclusion criteria, were recruited to participate in this study. Exclusion criteria included: 1) history of regular urinary incontinence; 2) treatment for or suspicion of prior toxic shock syndrome; 3) candidiasis; 4) active urinary tract infections; 5) active STIs; and 6) cervicovaginal irritation at the time of screening. Only asymptomatic women were included. Clinicians do not test for STIs in the asymptomatic population as these are not routine. We selected these exclusion criteria based on the indications from the Manual of Procedures of the Human Microbiota Project protocol [67]. However, we included women who took antibiotics during the last 2 months, since they were few (7.1%) and were distributed similarly among the women groups.

The date range in which human subjects’ data/samples were collected was between November 2017 and February 2020. The study and its procedures were conducted from March 2020 to September 2022. The study was approved by the Ethics Committees of the UPR-Medical Sciences Campus IRB (Protocol ref. 1050114/June 2014), San Juan City Hospital and has biosafety approval protocol # 94620. All subjects were informed (both verbally and in writing) of the sampling procedure, risks, and benefits of the study, gave written informed consent and signed HIPAA forms, following the Declaration of Helsinki. All staff involved in the project were certified by: CITI RCR, Social and Behavioral Research Best Practices for Clinical Research, HIPAA certifications and the NIH training on Protection of Human Subjects. Patients completed a metadata questionnaire with demographic characteristics (age, place of birth, employment, educational attainment), assessment of sexual risk (including the age of onset, current sexual partners), health history, antibiotic use, vitamins, and BMI.

A total of 367 samples were collected from which 337 where successfully sequenced and after rarefaction 4 samples were removed for low sequence count, for a total of 333 samples which corresponds to 294 women. Samples were collected using sterile Catch-All™ Specimen Collection Swabs (Epicentre Biotechnologies, Madison, WI), and placed in MoBio bead tubes with buffer (MoBio PowerSoil™ kit, MoBio, Carlsbad, CA) [67]. Swabs were then swirled for ∼20 seconds in 750 μL of MoBio buffer in the labeled specimen collection tube. For the sampling, a speculum was inserted for access and visualization of the cervix. The sterile swab was placed in the posterior fornix (cervix) and rotated along the lumen with a circular motion and swabs were immediately placed in the MoBio tubes. The area of the posterior fornix was used as a sample site to improve the detection of HPV and as a reliable indicator of the overall cervicovaginal environment, which has been demonstrated in previous research. [68]. Besides swabs, we collected approximately 10mL of cervical lavages by injecting PCR-grade sterile water into the vaginal canal for future studies. Collected lavage was transferred to a clean 15mL collection tube and pH was measured with hydrion wide range pH paper strips. All samples were coded and placed in ice up to 4 h, then samples were transported to the laboratory and stored at -80^0^C until nucleic acid extraction and PCRs, which was performed at a single laboratory (FGV) to reduce processing variation.

### DNA extraction and 16S rRNA sequencing

Genomic DNA extraction was done on posterior fornix (cervical) swabs using Qiagen Power Soil Kit (QIAGEN LLC, Germantown Road, Maryland, USA). No human DNA sequence depletion or enrichment of microbial or viral DNA was performed. A detailed description of the optimized extraction protocol is available in a previously published study [16]. In short, Powerbead tubes were homogenized for 10 minutes at 3,200 rpm, using the Vortex-Genie 2, G560 (Scientific industries, Inc. NY). We combined 100 μl of solution C2 and 100uL of solution C3 and vortexed for 5 seconds for cell lysis and, the elution solution was 100ul sterile PCR water, that was warmed to 55^0^C, and to increase DNA yield, allowed to remain on the filter for 5 min before the final centrifugation step. DNA concentration was measured using the Qubit® dsDNA H.S. (High Sensitivity) Assay (Waltham, Massachusetts, U.S.) (ranging from 5-100ng/μl). Genomic DNA from samples was shipped to an outsourced sequencing facility with an average genomic DNA concentration of 10-30ng/µL. No human DNA sequence depletion or enrichment of microbial DNA was performed.

The DNA obtained from cervical samples was normalized to 4nM during 16S rRNA gene library preparation. Universal bacterial primers: 515F (5’GTGCCAGCMGCCGCGGTAA3’) and 806R (5’GGACTACHVGGGTWTCTAAT3’) as in the Earth Microbiota Project (http://www.earthmicrobiota.org/emp-standard-protocols/16s/), were used to amplify the hypervariable V4 region of the 16S ribosomal RNA marker gene (∼291bp) [69]. The methodology applied for this process was identical to the one implemented in previous reports [16, 70-72]. Amplicons were quantified using PicoGreen (Invitrogen) and a plate reader (Infinite? 200 PRO, Tecan). Once quantified, volumes of each of the products were pooled into a single tube so that each amplicon is represented in equimolar amounts. This pool was then cleaned up using AMPure XP Beads (Beckman Coulter), and quantified using a fluorometer (Qubit, Invitrogen). Customized sequencing was outsourced at Argonne National Laboratory (Illinois, USA) using llumina MiSeq with the 2×250 bp paired-end sequencing kit. Amplicons were sequenced with the MiSeq Illumina platform in 3 different runs. The facility adds a negative control, and nothing is reported if they don’t produce sequence reads above 500 total reads. Internal positive controls are analyzed and aligned to the sequencer in real-time.

The reads obtained from the 16S-rRNA sequencing and corresponding metadata were uploaded in QIITA [73] study ID 12871 (https://qiita.ucsd.edu/study/description/12871). In addition, the resulting raw sequences were made available at the European Nucleotide Archive Project (ENA) under the access number ERP136546.

### HPV Genotyping and cytologic diagnoses

The kit used to complete the HPV genotyping consists of a highly sensitive short-polymerase chain reaction-fragment assay (Labo Biomedical Products, Rijswijk, The Netherlands, licensed Innogenetics technology). This process takes advantage of SPF10 primers to amplify a 65-bp fragment contained at the L1 open reading frame of HPV genotypes, followed by a Reverse-Hybridization reaction to determine which HPV genotypes are present in the sample by comparing the results to standardized controls provided by the kit. This assay facilitates the identification of the following mucosal HPV genotypes: 6, 11, 16, 18, 31, 33, 34, 35, 39, 40, 42, 43, 44, 45, 51, 52, 53, 54, 56, 58, 59, 66, 68, 73, 70, and 74 and classified as either high-risk (HR-HPV) or low-risk (LR-HPV). This methodology is explained thoroughly in a previously published study [16]. HPV genotype groups categorized samples as 1) only HR-HPV (exclusively HR types), 2) only LR-HPV exclusively low-risk HPVs, 3) Both (if a sample contained both HR and low-risk HPV genotypes, 4) HPV negatives or 5) missing (no HPV genotyping done on these samples) (**Table 1**).

Data from the medical records and questionnaires were obtained at time of patient visit and reviewed for results of cervical cytology and pathology reports. Data regarding abnormal cervical cytology was classified according to Bethesda system [74] in: ASCUS (atypical squamous cells of undetermined significance), ASCH-H (atypical squamous cells, cannot exclude high grade lesion), LGSIL (low grade squamous intraepithelial lesion), HGSIL (high grade intraepithelial lesion), or Squamous cell carcinoma (SCC). Missing values in the different categories corresponded to the fact that no information for a given category was retrieved for that specific participant.

### Sequence processing

Three Illumina demultiplexed paired-end sequence runs were uploaded and ran independently in QIIME2. DADA2 algorithm was used for truncating sequences based on quality, merging sequences, amplicon error correction, chimera identification, and obtaining representative sequences (features). Sequence length truncation was performed separately for each run, just before the drop of the base qualities. The maximum length truncation was 15 bases (reverse sequence). Taxonomy classification was performed using a Naïve Bayes classifier implemented in the q2-feature-classifier plugin [75] against a custom Greengenes modified database that contains sequences from Greengenes, the Human Oral Microbiota Database, and cervicovaginal vaginal reference sequences from the Human Microbiome Database [76] previously used [77-79]. Feature tables for the three runs were finally merged. Samples showed and average of 26,308 sequences/sample [min. 187 sequences, max. 80,253 sequences]. Rarefaction was run at 1,773 sequences per sample to include most of the samples, and four samples with a lower sequence number were eliminated (**Table S1**). All analyses were run with the rarefied table at the species level. An additional rarefaction of 5,000 sequences/sample was also run and feature tables were analyzed in parallel. For this threshold, a total of 16 samples were lost. Results between the two rarefaction cutoffs were similar. We also re-analyzed our data using the SILVA 138 database following the exact same parameters and metrics (see figures S6 and S7), also with similar results. The effect of the “sequencing run” was also included in all models of the analysis to account for the possible batch variability, however, no significant differences was observed.

### Statistical analysis

Population description was performed comparing the three main groups, non-pregnant, pregnant, and menopausal women. Categorical variables were compared with Fisher’s exact test and continuous variables with the Kruskal-Wallis test.

Cytology smears performed cervical lesion diagnosis, however, some women required biopsy. Therefore, a consensus result was used for the analyses. Only one sample was diagnosed as Squamous Cell Carcinoma (SCC) and it was included as HGSIL to facilitate the analysis. Additionally, diagnosis with ASCUS (5.4%, 14/258) was top-coded as LGSIL if they were HPV positive and negative when HPV negative. Then, the variable “Type of cervical lesions” only included LGSIL, HGSIL, and negative samples.

Comparison among HPV genotype categories explored first the distribution of women with the presences of any HR-HPV, but also having only LR-HPV genotypes, and then the distribution of women infected exclusively with HR-HPV or LR-HPV or mixed infections, meaning the presences of HR- and LR-HPV genotypes (called “both” on **Table 1** and **2**). Another comparison was also performed for mixed infections, although this one includes “Number of HR-HPV genotypes” or “Number of LR-HPV genotypes”, referring to the actual number of different HPV genotypes detected in a particular sample.

### Beta diversity

Beta diversity was evaluated using unweighted, generalized, or weighted Unifrac distances. Collinearity among variables was checked with spearman correlation using “cor.test” R function before including variables to the model. These included the following:

*distance ∼ Group (non-pregnant/pregnant/menopause) + Type of cervical lesion (LGSIL/HGSIL/negative) + HPV genotype (only HR-HPV/ only LR-HPV/ both HPV types/ negative) + BMI (numerical variable) + Antibiotics last 2 months (yes/no) + Sequencing run (Run A/B/C)*.

The trimester of pregnancy was also included only among pregnant women. For the stratified analysis for each of the women groups Age variable was also included. The variable “Sequencing run” was set as a random variable to account for the variability among runs.

Variables included in the model were analyzed using non-parametric Permutational Multivariate Analysis of Variance (PERMANOVA [80]) with “adonis2” function fr”m “ve”an” R package [81]. Permutation was set to 1000 and seed was set to 711. PERMANOVA model allows comparing variance between groups to the variance within groups (spatial location differences). “Run” was set as a random variable using “setBlocks” function from the “permute” R package [82].

Pairwise analyses for the significant categorical variables were performed with “pairwise.adonis2” function from the “pairwiseAdonis” R package (Martinez Arbizu 2017).

### Taxa association with study variables

Microbial taxa and their association with all the above variables were assessed using Maaslin2’s linear model in R [83], we also set “sequencing run” as random variable. Maaslin2 is a bioinformatic tool that helps to identify taxa-variables associations. The output table (**Table S3**) shows the specific taxa associated with a particular variable. The table also provides the model coefficient value (effect size), the P-value, and the P-value adjusted for false discovery rate (“fdr”), among other parameters. If the variable is categorical, a reference is needed; negative samples were always set as the reference for our analysis. A positive coefficient indicates a positive association (increase in a taxon abundance) with a particular variable. A heatmap is also generated where the plus sign in the red squares suggests an increase in the taxa abundance for a specific variable, while a negative sign in the blue squares indicates a decrease of abundance, always based on the reference (**Fig. 1-E**). The model included women’s group, HPV types, type of cervical lesions, and significant variables in the beta diversity analysis (pH, age).

### Alpha diversity

Microbial alpha diversity was measured with the Shannon index calculated using “diversity” function from “vegan” R package [81] and the Chao1 estimator calculated using the “apply” function from the “OTUtable” library [84]. Shannon index provides information about richness and evenness of a microbial community by sample, while Chao1 estimator provides only the microbial richness estimated in an exhaustive sampling scenario. Linear mixed-effect model (LMM), “lmer” function from “lmerTest” R package [85] were fitted, setting “Sequencing run” as the random variable to account for the differences in sequencing facilities. Fixed (explanatory) variables included were the same that for the beta diversity analysis. To obtain the best-fitted model we run the “step” function from “lmerTest” R package [85].

The trimester of pregnancy was also included only among pregnant women. For the stratified analysis for each of the women groups “age” variable was also included. Collinearity among variables was checked with spearman correlation using “cor.test” R function, before including variables to the model. We also checked the normality of the linear regression residuals and did logarithmic transformation to alpha diversity metrics to reach residual normal distribution when necessary. Models were run for between 164 and 184 samples due to missing samples. R library “emmeans” [86] was used to run pairwise analyses for mixed model.

### Community State Types (CSTs)

Each women’s sample was classified into CST following the protocol of Valencia program [18]in Python 3 (https://github.com/ravel-lab/VALENCIA). Input data had to be formatted using local scripts. To compare the proportion of CST in women in Puerto Rico from this study, with a similar Hispanic population but living in the USA, metadata from a very known study of cervicovaginal microbiota were downloaded and compared [12]. Prevalence of CSTs was also compared among women groups and among cervical lesions. A 95% confidence interval for these values were calculated using “BinomCI” function from “DescTools” R package [87].

### Longitudinal changes of the cervicovaginal microbiota in a subset of women

To explore whether microbial profiles in this population vary over time we analyzed two-time points longitudinal data. Thirty-nine women who had had a second sampling between 4 to 16 months after their first visit were analyzed. We first used the CST classification to group women with CSTs dominated by *Lactobacillus* (CST-I, II, III and V) or diverse (CST-IV). We then built a table to observe the coincidences first among type of cervical lesions and then among positive and negative HPV infection, finally we ran a Fisher’s exact test for each table (**Table S7** and **S8**).

As supplementary materials we provide the updated STORMS checklist (The Organization and Reporting of Microbiome Studies’) which we updated based on our study’s criteria to provide complete reporting of our study and ensure reproducibility.

## Acknowledgments

We extend our heartfelt gratitude to the volunteer students who took part in this study. Additionally, we acknowledge Rafael López Martinez building scripts for data analysis.

## Author contributions

Conceptualization: FGV

Data curation: ADM, J.R., DVR, FGV Formal analysis: DVR, FGV

Funding acquisition: FGV Investigation: FGV, J.R.

Methodology: FGV, J.R., IAV, ETR, ADM, L.J.F., MS, MMF

Project administration: FGV Supervision: FGV, J.R.

Writing – original draft: DVR, FGV

Writing – review & editing: FGV, J.R., IAV, ETR, JAG, KJW, ADM, L.J.F., MS, MMF

## Data availability statement

Data is available. The reads obtained from the 16S-rRNA sequencing and its corresponding metadata were uploaded in QIITA [73] study ID 12871 (https://qiita.ucsd.edu/study/description/12871). In addition, the resulting raw sequences were made available at the European Nucleotide Archive Project (ENA) under the access number ERP136546 https://www.ebi.ac.uk/ena/browser/view/PRJEB51893?show=reads.

Supplement data is also available.

## Funding

Puerto Rico Science and Technology and Research Trust award, #2020-00112 Center for Collaborative Research in Minority Health and Health Disparities (RCMI), 2U54MD007600

The Hispanic Alliance for Clinical and Translational Research (Alliance), U54GM133807 NIGMS RISE Program at the UPR Medical Sciences Campus, R25GM061838 Hispanics-In-Research Capability: SoHP & SoM Partnership (HiREC), S21MD001830 The Puerto Rico IDeA Networks of Biomedical Research Excellence; Advancing Competitive Biomedical Research in Puerto Rico, 5P20GM103475-20 Center for Interdisciplinary Research on AIDS, Yale University, P30MH062294

